# ATR-FTIR spectra of polysaccharide isolated from *Euphorbia caducifolia* and its chemical modifications: A study of Principal component analysis

**DOI:** 10.1101/2023.03.10.532019

**Authors:** Kusuma Venumadhav, Kottapalli Seshagirirao

## Abstract

Polysaccharides are the carbohydrates composed of same or different types of saccharides in an organized manner with glycosidic bonds. These are useful for various industrial applications mainly in food and pharmaceutical companies. Modifications of the polysaccharides change its structure and lead to the physical as well as functional differences. Mostly, chemical methods are used to modify its structure to study the structure-activity relationship in interest towards its physical properties like solubility and biological activities among various types of molecular modifications. In this study, a polysaccharide isolated from stems of *Euphorbia caducifolia* and modified its structure chemically by substituting different groups by sulfonation, carboxymethylation, acetylation and phosphorylation with various degree of substitutions. Molecular modifications of the polysaccharides showed the impact on water solubility as a physiochemical property. Thus, ATR-FTIR spectroscopy was employed along with a chemometric analysis, i.e., Principal components analysis (PCA) to understand and identify the chemical moieties differed from the unmodified polysaccharide. From PCA analysis, PC_1_ and PC_2_ components suffice to differentiate the modified polysaccharides with its native form.

## Introduction

Polysaccharides are the carbohydrates composed of same or different types of saccharides in an organized manner with glycosidic bonds. These are used for various industrial applications mainly in food and pharmaceutical companies. Modifications of the polysaccharides change its structure and lead to the physical as well as functional differences (Cumpstey, 2013). Mostly, chemical methods are used to modify its structure to study the structure-activity relationship in interest towards its physical properties like solubility and biological activities among various types of molecular modifications. Polysaccharide hydroxyl groups are major contributors for chemical modifications. Apart, Alginates and pectins are composed of carboxylic groups called uronic acids. In modifications, Saccharide hydroxyl oxygen act as nucleophile and carbon serve as an electrophile. Nucleophile oxygen can undergo chemical reactions like Etherification (carboxymethyl ether, hydroxyethyl ethers), Esterification (acetyl, carboxyl esters, sulphonate esters) and electrophile carbon can undergo introducing the groups like sulfate and halide groups (Cumpstey, 2013). Additionally, oxidation of sugars, nitrogen group reactions of amino sugars, esterification and other substitutions on carboxylic groups of uronic acid sugars are also considered. All together gives change in chemical, physical and biological changes.

## Materials and Methods

### Sulfonation

50 mg of polysaccharide dissolved in 10 ml of formamide and 10 ml of pyridine mixture. After, added 4 ml of chlorosulfonic acid on ice bath about 2 hrs. Placed the reaction mixture for 15 hrs at 4°C and ended the reaction by adding cold water. Sodium hydrogen carbonate used to neutralize the reaction mixture until it ceases the effervescence. Then, dialyzed the solution about 7 days against distilled water and concentrated under reduced pressure and lyophilized (O Neill, 1956).

### Carboxymethylation

Yang et al. 2011 method was followed with minor modifications (Yang et al., 2011). 10 mg of polysaccharide suspended in 4 mL of isopropanol and kept under stirring at RT for 15 min, followed by the slow addition of 1.5 mL of 20% NaOH with 1 h stirring. Then the solution of 150 mg/ ml of chloroacetic acid was added to the reaction mixture with stepwise under constant stirring. The reaction was continued for 4 h at 60°C. Cooled down to RT, and 0.5 M of hydrochloric acid was added to neutralize the mixture. The derived polysaccharide was dialyzed against normal water for 6 h and then with distilled water for 12 h. Dialysate was precipitated with ethanol and lyophilized.

### Acetylation

50 mg of ECP dissolved in 7 ml of formamide and further added 12.5 ml of pyridine. After, added 10 ml of acetic anhydride at 4°C about 2 h at frequent intervals. The reaction was continued for 20 h at RT and ceased the reaction by adding cold water. Sodium hydrogen carbonate used to neutralize the reaction mixture until it ceases the effervescence. Then, dialyzed the solution about 7 days against distilled water and concentrated under reduced pressure and lyophilized (Mellerowicz, 2013)

### Phosphorylation

50 mg of polysaccharide was suspended in 10 ml of N, N-Dimethylformamide (DMF) along with 4 g of urea and stirred for 1 h to homogenize the mixture. 50 mg of phosphoric acid was added and stirred for 4 h at 130°C. Reaction mixture was brought to RT, filtered, and washed with a mixture of 1-propanol, and distilled water followed by 0.1mM hydrochloric acid, and with distilled water. The obtained filtrate was lyophilized (Oshima et al., 2008).

### Morphological and elemental changes

Morphological changes of the modified polysaccharides were studied under scanning electron microscopy (SEM), and the change in the elemental composition was investigated with energy dispersive x-ray spectroscopy (EDS).

### Functional group studies and PCA analysis

Chemical modifications of the polysaccharide further confirmed with functional group analysis by IR spectroscopy. Minor changes in the chemical modifications and major distributions of the different functional groups of the modified polysaccharides were studied under the statistical analysis of principal components. PC1 and PC2 were considered for the analysis (Szymanska-Chargot & Zdunek, 2013).

### Biological significance

Change in the chemical structure always changes its biological activity. Antioxidant activity and cytotoxicity of the modified polysaccharides were studies with previous methods.

## Results and discussions

### Morphological and elemental analysis

Modified polysaccharides changed their morphology as they introduced by other functional groups. Sulfated, carboxymethylated polysaccharide looks like fabric like structure and flakes like structures are observed with acetylated and phosphorylated polysaccharides whereas the native form of ECP is crystalline like structure (Fig.3.2.1). Change in the elemental composition of the carbon and oxygen of the native form by introducing acetyl and carboxymethyl groups are not much significant. However, the important observation can make from the composition that, decrease in the “C” element from native 51% to 48% and 45% of acetyl and carboxymethyl respectively propose the etherification of the polysaccharides. 8% of the sulfur element and 4% of phosphorous elements confirm the substitution of sulfate groups and phosphate groups on the polysaccharide (Fig.3.2.2).

**Fig.3.2.1.**
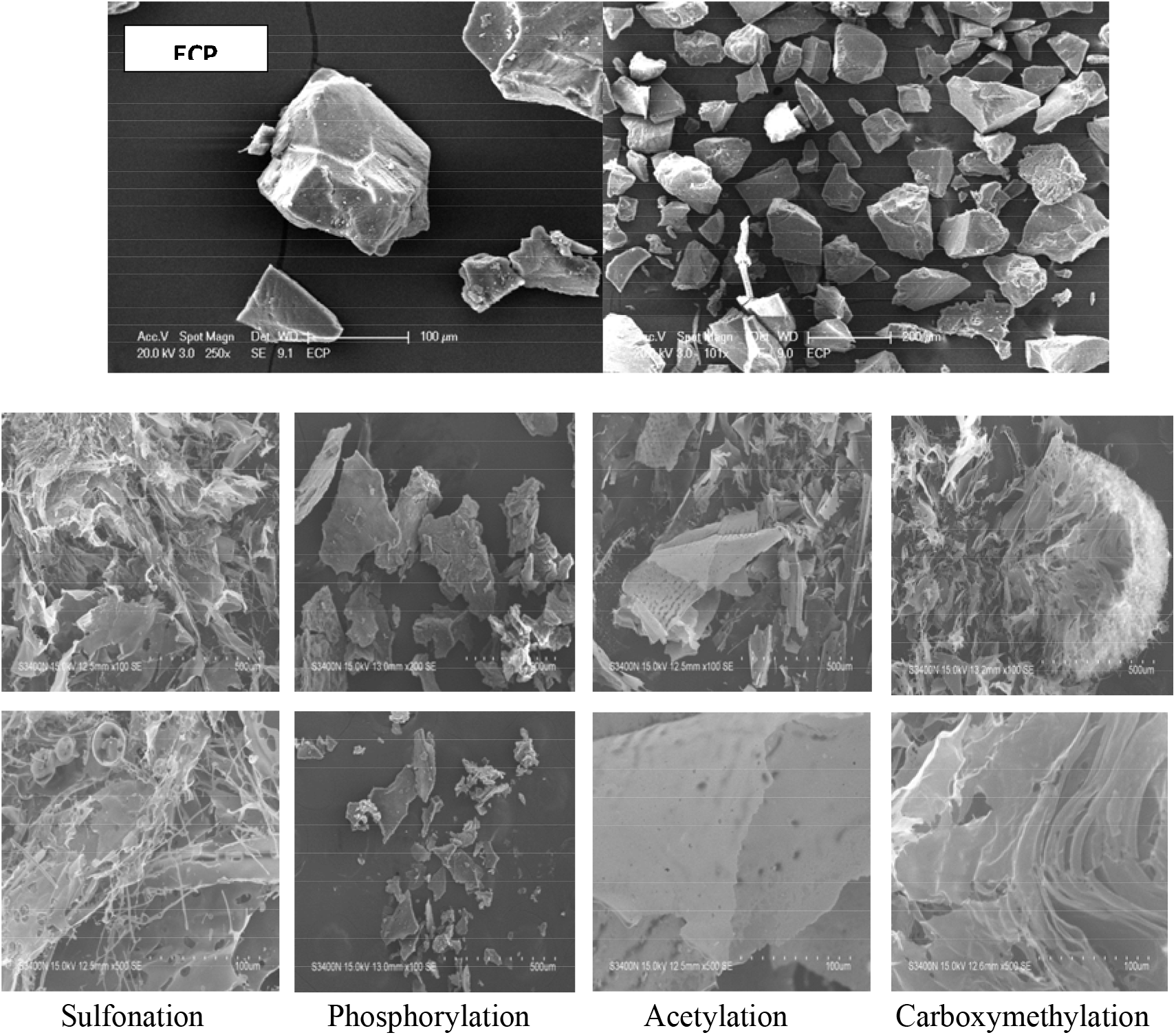
Scanning Electron Microscopy images of native and modified polysaccharides.

**Fig.3.2.2.**
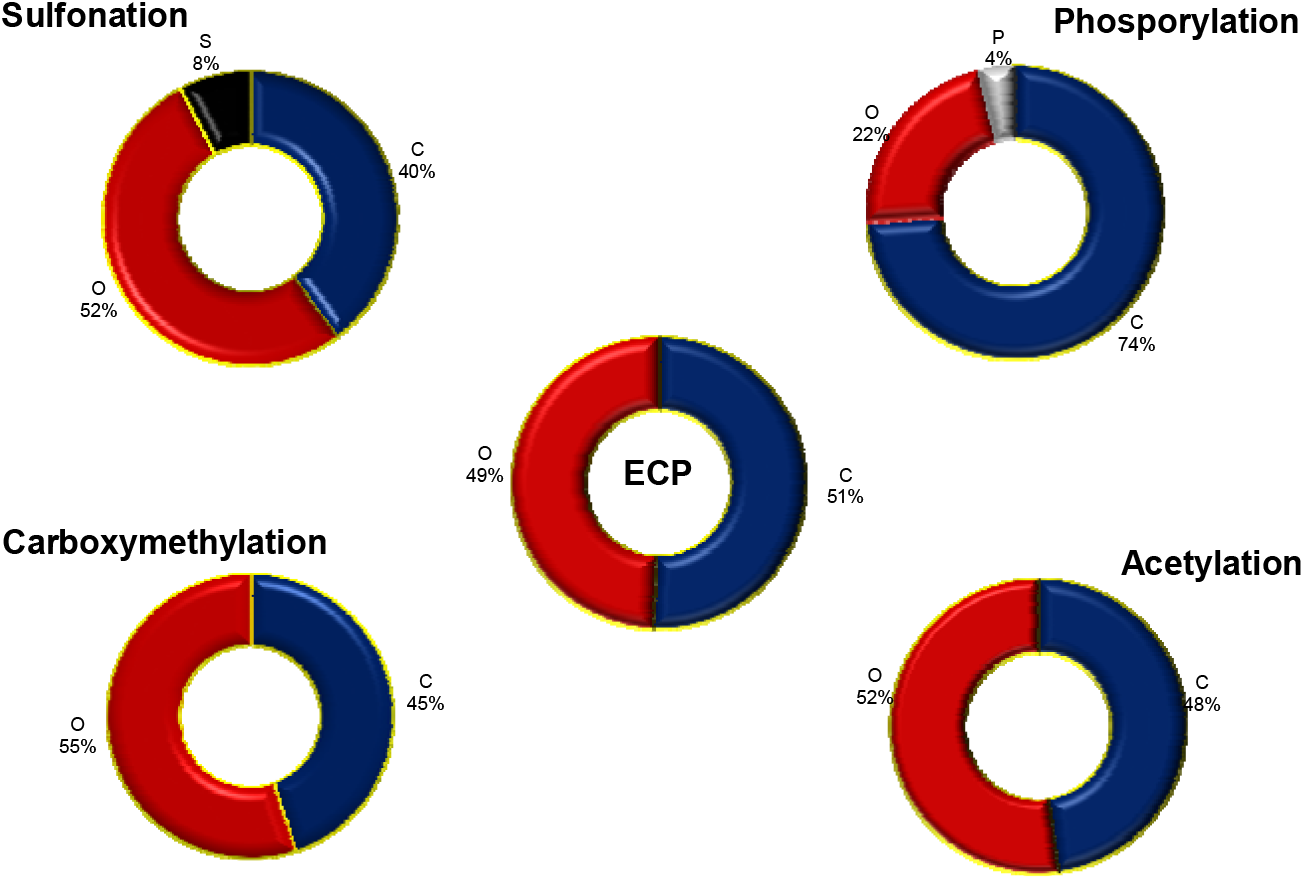
Elemental composition of native and modified polysaccharide.

### Functional group studies and PCA analysis

An IR spectrum of the modified polysaccharides confirms the presence of relative functional groups respective to the modifications. Carrageenan, a natural sulfated polysaccharide obtained from marine algae. Sulfated polysaccharides are familiar with many biological activities like anti-tumour, antithrombin, anti-viral and mostly for anticoagulant activity (Yuan et al., 2005). Heparin know to be a natural anticoagulant, activity depends on the degree of sulphation of heparin (O Neill, 1956). A peak at 1260-1210 cm^-1^ confirms the sulphation of the polysaccharide along with sugar sulfate peals around 800 cm^-1^ (Gomez-Ordonez & Ruperez, 2011). Phosphate of the polysaccharide gives P=O, P-O stretch vibrations at 1200-1100 cm-1. For carboxymethylation and acetylation, CO stretches at 1900-1550 cm^-1^ due to the carbonyl group and stretch at 1800-1740 cm^-1^ assigned to carboxylic groups. Similar stretches were noticed with respectively modified polysaccharide in comparison to native form (Fig.3.2.3). Change in the intensity of OH stretch of alcoholic hydroxyl groups confirms the derivative forms of esters.

**Fig.3.2.3.**
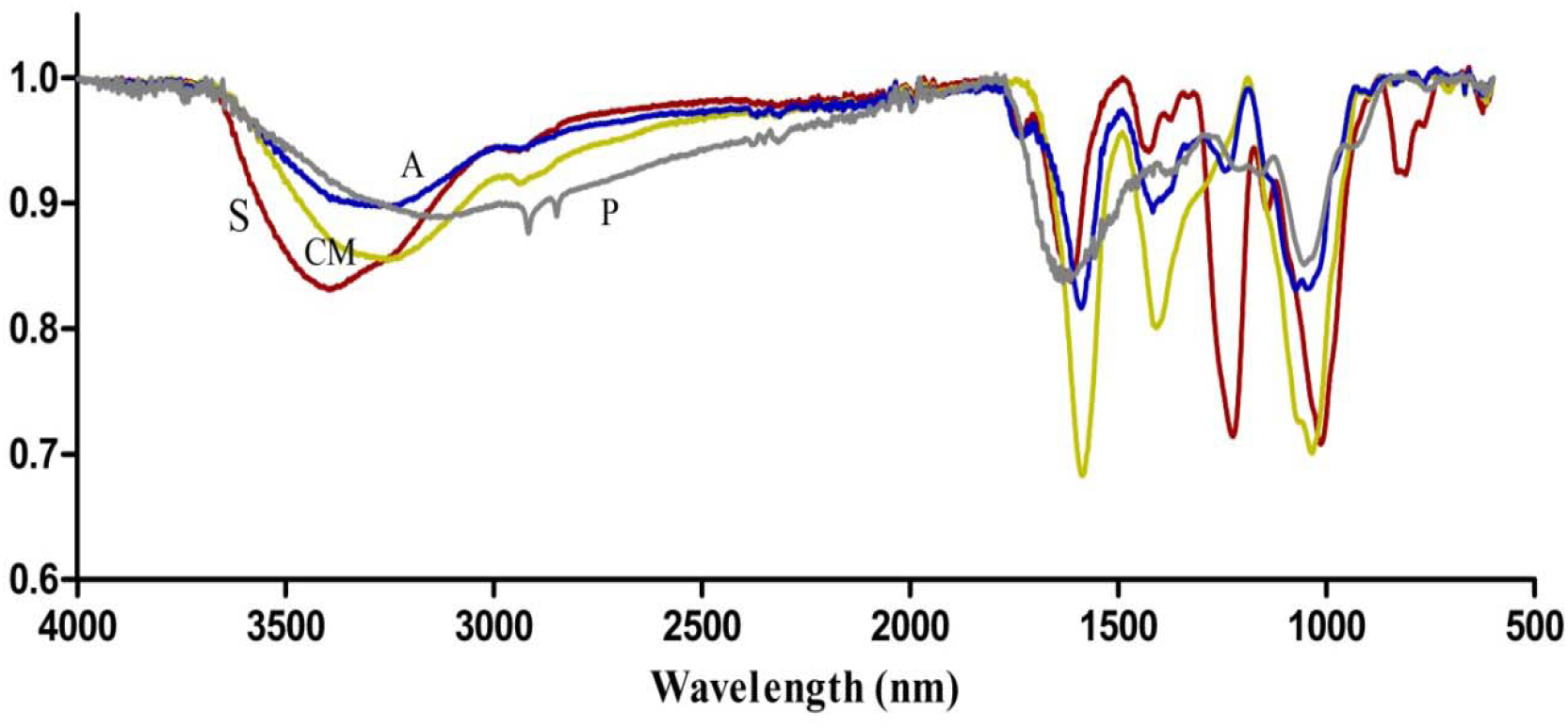
ATR spectra of modified polysaccharides; A-Acetylated, CM-Carboxymethylated, S-Sulphated, P-Phosphorylated polysaccharides.

Multiple variation analysis is used to analyze the large set of data to limited variable interpretation like IR spectral data. PCA is one of the methods used to characterize the modified spectra from its native form. Total 75.68% of variation was observed with first 2 principal components PC_1_ and PC_2_ and considered to categorize the observations. A negative correlation was noticed with sulfated and phosphate forms whereas togetherness of native form with carboxymethylated and acetylated forms (Fig.3.2.4 A). Biplot shows the influence of variables on observations with particular stretch levels of vibrations at respective wavelengths (Fig.3.2.4 B). The combination of the various observations concerning their modifications alters the variation to the 79.94% from 75. 68%. (Fig.3.2.4 C).

**Fig.3.2.4.**
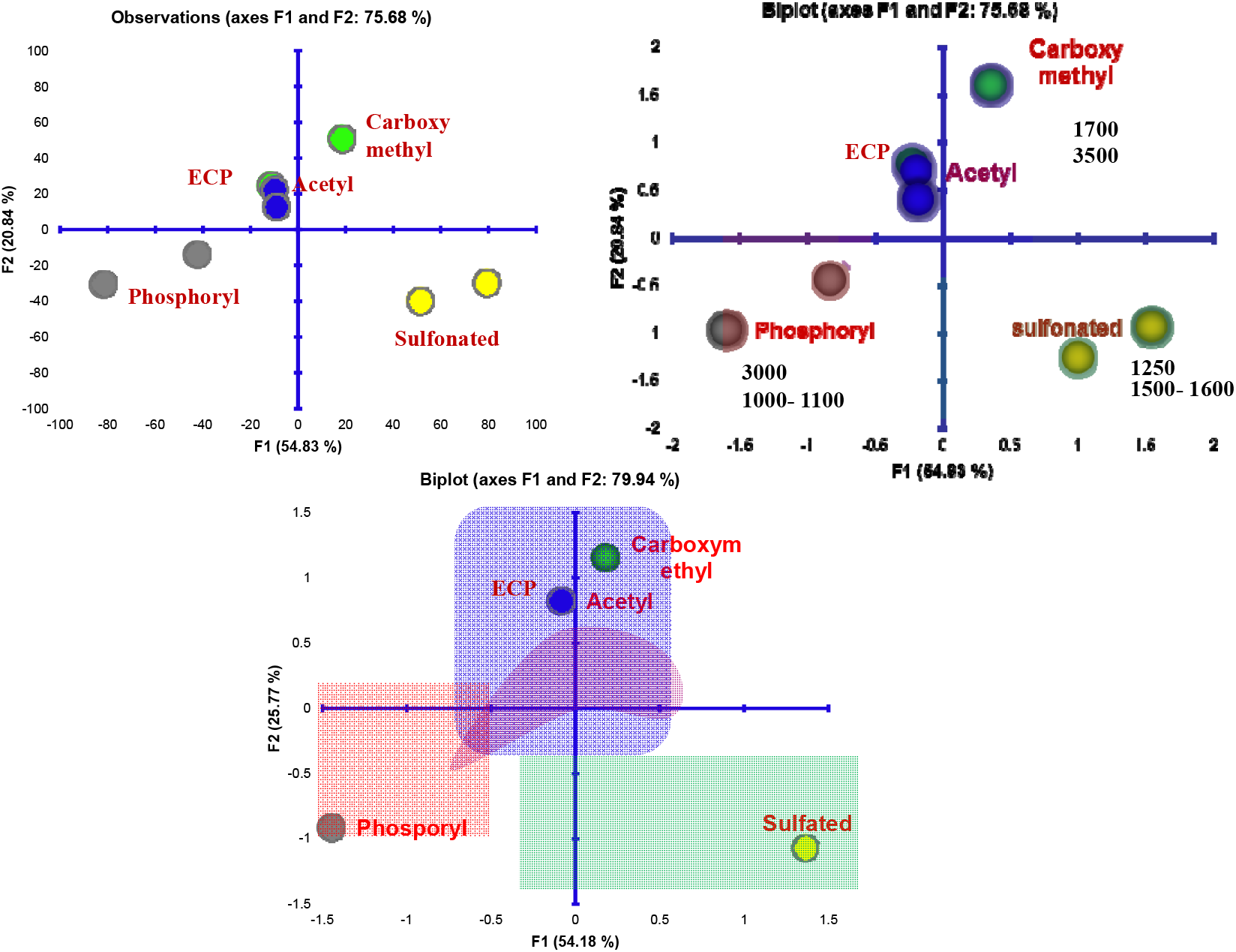
Principal Components analysis of modified polysaccharide spectra; A) Plot of PC_1_ and PC_2_ of observations, B) Plot of PC_1_ and PC_2_ observations with variables, C) Average plot of observations.

### Biological Significance

Structural changes always modify the biological activity. Structural changes employed to improve the physicochemical barriers like solubility, flow properties, and others. In addition, Biological significance also greatly mentioned. The addition of sulfate group improves the anticoagulant activity and antioxidant activity (O Neill, 1956). The ability of free radical scavenging activity of increased with the addition of carboxymethyl, sulfate and acetyl groups to the native form of polysaccharide. In contrast, phosphate introduced polysaccharide decreased its ability to scavenge the DPPH free radicals. A polysaccharide isolated from the plant does not show any toxicity towards macrophages. The same results were observed with carboxymethylated and acetylated ECP whereas cell death was noticed with sulfated and phosphorylated ECP.

**Fig.3.2.5.**
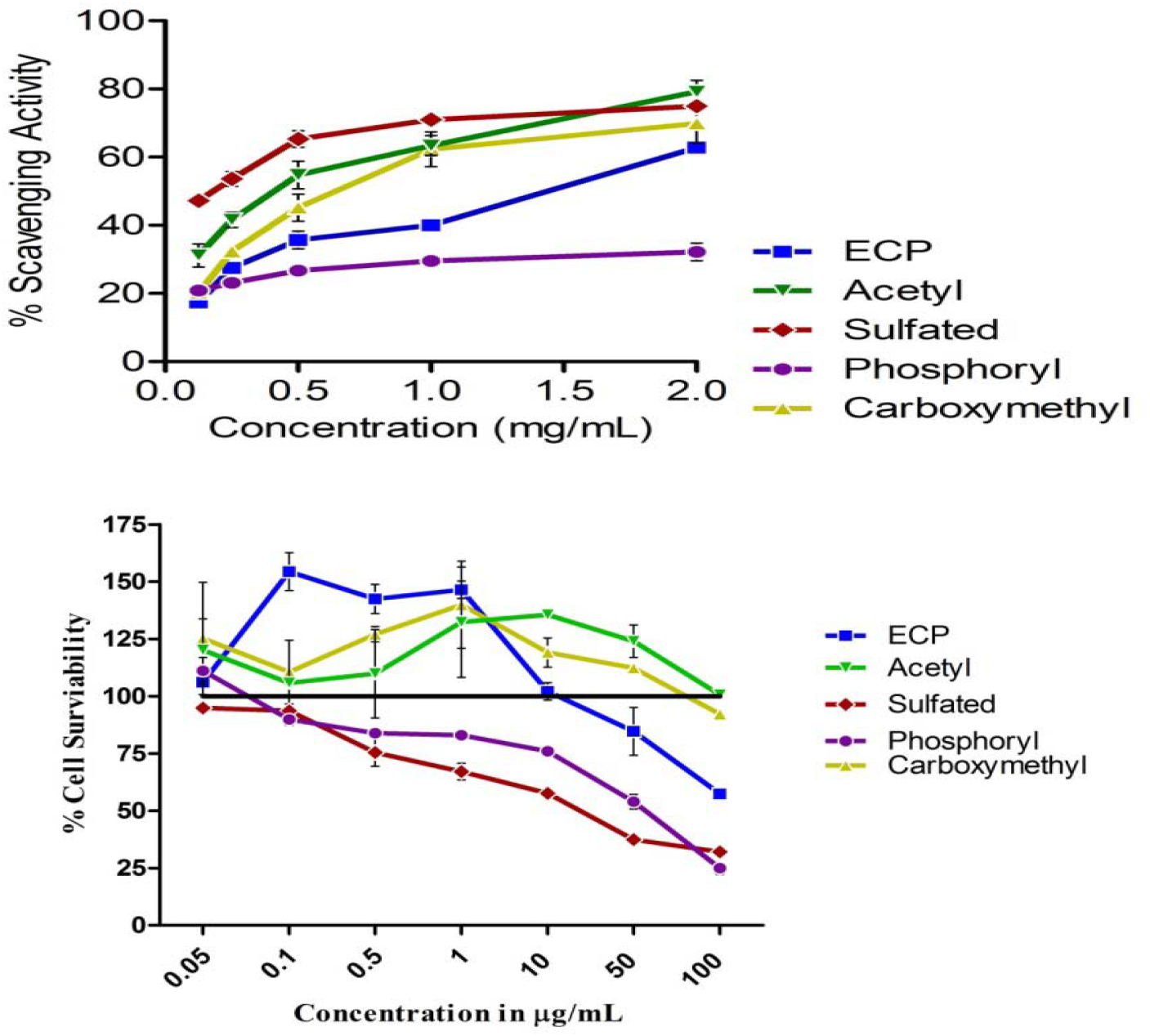
The biological significance of Modified polysaccharides: A) Antioxidant activity, B) Cytotoxic activity.

## Conclusions

Change in the structural aspects of the molecules alters their nature towards physicochemical and biological properties. 8% of sulfate and 4% of phosphate groups introduced. Change in the elemental composition of C and O gives the alteration carboxymethyl and acetyl polysaccharides. Principal components analysis of ATR spectra discriminate the respective polysaccharides each other and with native form. Biological significance greatly altered with modified polysaccharides to native form.

